# Where to stop Ψ-BLAST iterations so that the found sequences remain related to the query sequence?

**DOI:** 10.1101/2023.10.24.563694

**Authors:** Natalya S. Bogatyreva, Alexei V. Finkelstein, Dmitry N. Ivankov

## Abstract

The BLAST program [Altschul et al., 1990, J. Mol. Biol., 215:403-410] stands as a widely used tool for the search of the most similar sequences, while the iterative Ψ-BLAST program [Altschul et al., 1997, Nucl. Acids Res., 25:3389-3402] offers a high sensitivity for detecting remote homologs of the query sequence through an iterative usage of the BLAST search. However, the number of iterations that have to be used by the Ψ-BLAST is rather poorly justified in the literature. Our study shows that, as the number of iterations increases, Ψ-BLAST rapidly loses the ability to be guided by the query sequence in the search for homologs. When working with the non-redundant (nr) sequence database of 2021, Ψ-BLAST, already after the second iteration, retains the query sequence at the top of the list of the found homologs to this sequence in only 18% of cases. Moreover, a query sequence is still listed among homologs found by Ψ-BLAST after the recommended 10 iterations [Altschul et al., 1997, Nucl. Acids Res., 25:3389-3402] in only 42% of cases. Using a considerably smaller nr database-2011 as a reference, we reveal that these effects intensify over time. Our findings underscore the necessity for circumspection when interpreting Ψ-BLAST outcomes; the degree of vigilance must increase with the database size. A vigilant monitoring of the position of the query sequence in the array of detected homologs is needed. We recommend using the disappearance of the query sequence from the list of homologs produced by Ψ-BLAST as a criterion to conclude the Ψ-BLAST iterations.

## Introduction

The Basic Local Alignment Search Tool (BLAST) stands as the foremost program employed for sequence similarity searches [Altschul et al., 1990]. Primarily, it addresses two interconnected yet distinct objectives: (i) identification of homologs within a sequence database for a given query sequence and (ii) alignment of the found homologs with the query sequence. The BLAST’s widespread adoption is underscored by the remarkable fact that the BLAST paper [Altschul et al., 1990] holds a position among the most cited articles in the Journal of Molecular Biology, boasting 106,846 citations (as of August 6, 2023). The BLAST search serves as the workhorse and starting point for nearly every task in bioinformatics, spanning from protein 3D structure prediction pipelines [Martí-Renom et al., 2000] to the prediction of signal peptides [Frank, Sippl, 2008], protein function [Conesa et al., 2005], and other various attributes inferred from homology [Gardy et al., 2005; Moriya et al., 2007].

Position-Specific Iterated BLAST (PSI-BLAST or Ψ-BLAST) is an immensely popular tool purpose-built for increased sensitivity in the pursuit of remote homologs: if their structure or function is already known, they can hint at the structure or function of the target protein. The Ψ-BLAST paper [Altschul et al., 1997] stands as the most cited article in the Nucleic Acids Research journal, amassing an impressive 82,916 citations (as of August 6, 2023).

The first iteration of Ψ-BLAST represents a conventional BLAST search for homologs of the query sequence; but, starting from the second iteration, the set of homologs found in the preceding iteration guides the construction of a “sequence profile” used in the current iteration for the search of homologs of this sequence profile. This distinctive approach empowers Ψ-BLAST to find distant homologs that often remain beyond the reach of a typical BLAST search.

The maximum iteration count serves as an essential parameter in a Ψ-BLAST search. However, consensus remains elusive regarding the optimal number of iterations to employ. The default value in the stand-alone BLAST package is ten Ψ-BLAST iterations [Altschul et al., 1997]. Various researchers have conducted from 2 [Lu et al., 2004] and 3 [Cheng, Baldi, 2007; Chou, Shen, 2007] to 16 [Altschul et al., 1997] iterations. Increasing the maximum iteration count is thought to augment the search sensitivity by unearthing progressively more remote homologs. Hence, at first glance, the choice of the maximum iteration count seems to be just a question of a balance between extra search sensitivity and running time. However, if the query sequence is present in or inserted into the database of explored sequences, its position within the Ψ-BLAST-produced homolog list could change from iteration to iteration. While it is rational to expect the query sequence to lead the pool of sequences homologous to itself at each iteration, some query sequences lose the first position after a certain iteration. Ultimately, the query sequence may even slip out of the homolog list entirely.

In the present study, we systematically investigate the behavior of the standard Ψ-BLAST search with respect to its ability to keep focused on the query sequence. We demonstrate that Ψ-BLAST quickly deviates from the query sequence which is present in the explored database, and this phenomenon gets worse as the sequence database gets larger. In the non-redundant (nr) database-2011, a mere 36% of Ψ-BLAST searches retain the query sequence in the top position of the list of found homologs already after the second iteration. For the richer nr database-2021, this proportion diminishes to just 18%. Moreover, the query sequence disappears from the list of homologs before the Ψ-BLAST’s convergence in ≈26% of cases for the nr database-2011, and in ≈58% of cases for the nr database-2021. These observations are complemented by the fact that the standard (limited to 10 iterations) Ψ-BLAST search converges only in approximately half of cases for the nr database-2011, and only in around a quarter of cases for the nr database-2021. In light of these results, it is prudent to exercise caution when interpreting Ψ-BLAST outcomes. At each iteration, it is recommended to track the position of the query sequence in the list of found homologs and consider its disappearance from the first position of this list, and even more so from the list in general, as a criterion for halting Ψ-BLAST iterations.

## Results and Discussion

To systematically examine the behavior of query sequences in the Ψ-BLAST searches, it was imperative to assemble a substantial protein sequence pool to get robust statistics on the query sequence behavior. As such, we opted to randomly select 1000 sequences for our computer experiment — this number was deemed sufficient to ensure a diverse range of scenarios and to yield robust statistics.

With the aim of tracking the query sequence behavior over time, we focused on two arbitrary time points: the databases of years 2011 and 2021. A 10-year difference was considered ample to draw meaningful conclusions about long-term trends related to the investigated phenomenon. To ensure the presence of all query sequences in both databases, we extracted these sequences from the SwissProt database dated back to 2011 and made sure that in the database-2021 they are the same. For each query sequence selected, we executed 10 iterations of the Ψ-BLAST search, with default parameters. These searches were conducted against the non-redundant (nr) databases for 2011 (∼13,000,000 sequences) and 2021 (∼300,000,000 sequences), both provided within the Ψ-BLAST package [https://ftp.ncbi.nlm.nih.gov/blast/db/].

A typical instance of the Ψ-BLAST search is depicted in Figure 1. After the first iteration, the top spot is always occupied by the query taken from the database or a sequence that is 100% identical to it. However, merely after the second iteration, this query sequence descends to the 63rd position, followed by a further drop to the 83rd spot during the third iteration, where the Ψ-BLAST search has converged.

**Figure 1.**
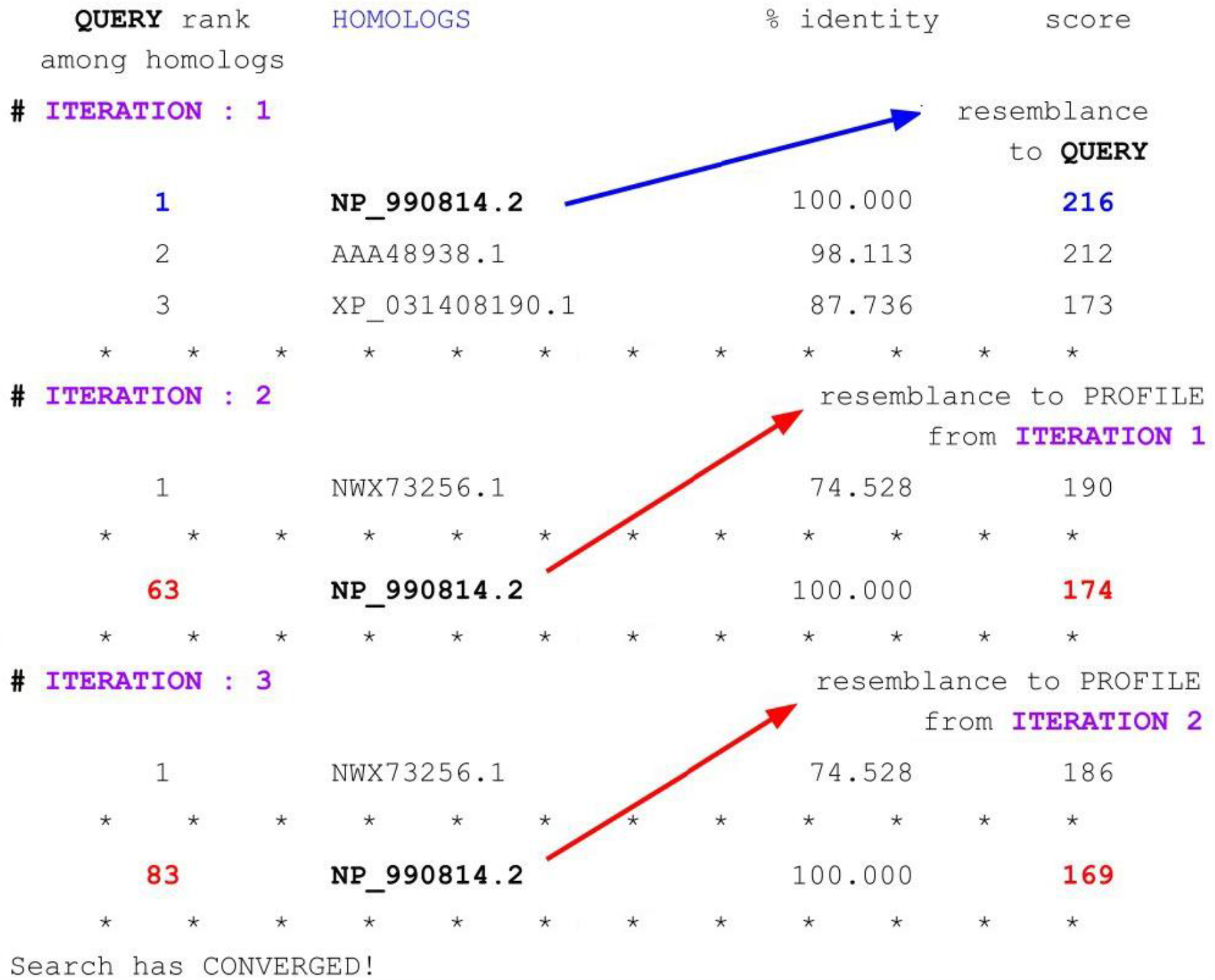
A typical Ψ-BLAST search. The blue arrow indicates that the sequence similarity to the QUERY sequence defines the score at the first iteration. Notably, the score between the QUERY sequence and itself (216, highlighted in blue) substantially surpasses the scores between this sequence and the PROFILEs at the following iterations (174 and 169, indicated in red within the score column). The red arrows at these iterations point to the exclusive importance of the score of the sequence similarity to the PROFILE built after the previous iteration.

The reason for the disappearance of the query sequence from the top position is that the Ψ-BLAST ranking is based on the similarity of the sequence namely to the PROFILE obtained from the previous iteration and not to the query sequence. As the iterations progress, the alignment score of the query sequence deteriorates. The dissimilarity in scores between the 1-st and the 2-nd, and between the 2-nd and the 3-rd iterations is attributed to the deviation of the sequence PROFILEs used at Iterations 2 and 3 from the query sequence. At Iteration 1, the “PROFILE 1” consisted of the QUERY sequence only; at Iteration 2, the “PROFILE 2” was the result of averaging over the 250 (by default) “most homologous to the PROFILE 1” sequences; at Iteration 3, the “PROFILE 3” was the result of averaging over the 250 “most homologous to the PROFILE 2” sequences.

Figure 2 portrays examples of all observed scenarios. A Ψ-BLAST search can converge after merely the second iteration, though even after the 10th one (the last, by default) it often does not converge. Following the second iteration, the search may reach convergence, with the query sequence occupying different positions: (i) the highest rank in the search, (ii) a position within the list of the default 250 homologous sequences intended for incorporation into the subsequent iteration’s PROFILE, or (iii) being excluded from the list of 250 identified homologs (represented by the “>250” symbol).

**Figure 2.**
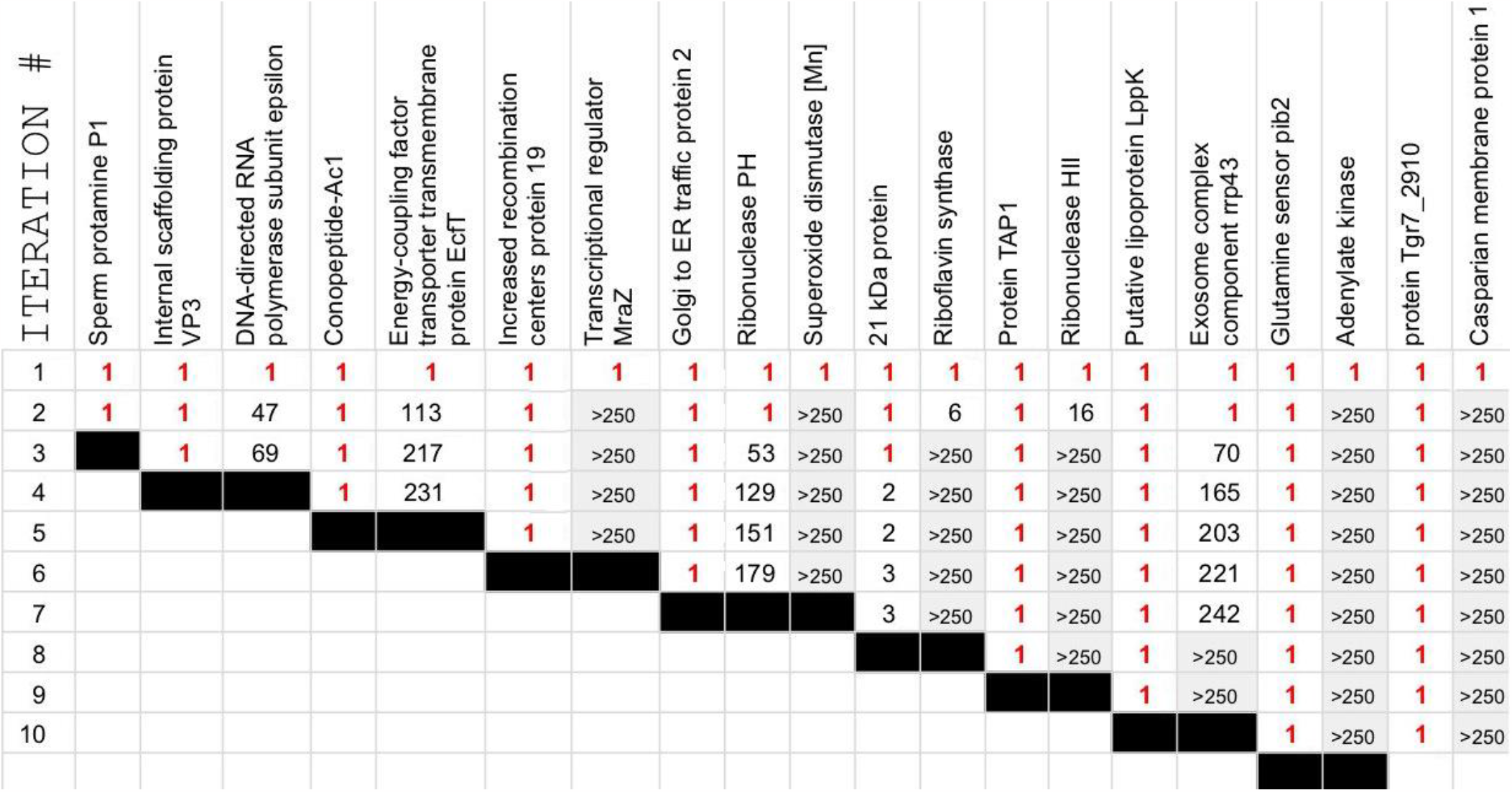
A collection of twenty diverse variants for the search progress conducted by Ψ-BLAST, encompassing the entire spectrum of observed scenarios. The query proteins are organized into columns with the protein name at the top; the leftist column designates the current iteration number. Other numbers denote the position of the query sequence in the sorted list of homologs resulting from the given Ψ-BLAST’s iteration. The place 1 in the homologs list is highlighted in red. The symbol “>250” on the gray background denotes the exclusion of the query sequence from the list of the default 250 “most homologous to the PROFILE used at the given iteration” sequences (these 250 sequences are to be incorporated into the next-iteration PROFILE). Convergence of the Ψ-BLAST search is depicted by a black rectangle.

Among these scenarios, the prospect that the query sequence can be washed out from the list of homologous sequences is the most alarming. In this case, given that the PROFILE for the subsequent iteration is constructed from the found 250 “best” homologs, the query sequence ceases to contribute to the formulation of the PROFILE for ensuing iterations. Consequently, commencing from the iteration marked by the “>250” symbol in Fig. 2, the query sequence no longer steers the trajectory of the Ψ-BLAST search during all the subsequent iterations. This particular situation can materialize surprisingly early — after the second iteration. The proteins “Transcriptional regulator MraZ”, “Superoxide dismutase [Mn]”, “Adenylate kinase”, and “Casparian membrane protein 1” in Fig. 2 feature prominently in the described scenario with additional 3, 4, 8, and 8+ iterations, respectively, before achieving convergence.

To attain a clearer comprehension of the “washing out of the query sequence” phenomenon, we provide schematic representations of the Ψ-BLAST searches within the sequence space for the “DNA-directed RNA polymerase subunit epsilon” (Fig. 3A) and the “Ribonuclease HII” (Fig. 3B) proteins, both being featured among the examples showcased in Fig. 2. The region of homologs discovered in the next iteration may continue to encompass the query sequence’s protein family, but it also may abandon some part of that family, including sometimes (Fig. 3B) even the query sequence itself. This disappearance of a query sequence occurs if the iterative search shifts to regions that contain more and more sequences.

**Figure 3.**
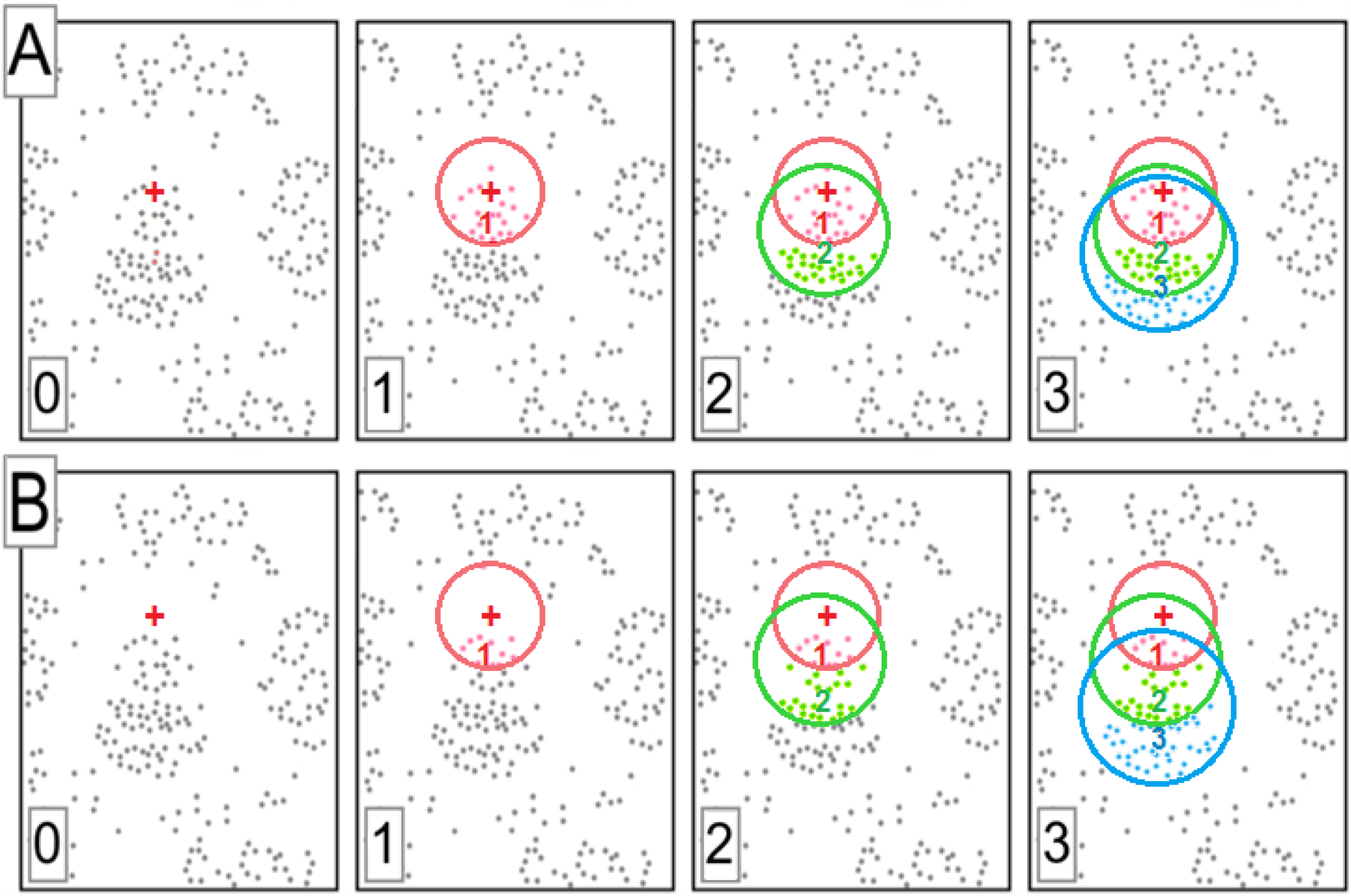
A schematic presentation of two distinct scenarios delineating the course of a Ψ-BLAST search within the sequence space over the initial three iterations. Each image captures the situation at discrete moments of the search: start position (0), and positions after the subsequent first (1), second (2), and third (3) iterations. Protein sequences are symbolized by dots, while the query sequence is marked by a red ‘**+**’ symbol. Red, green, and cyan circles delineate the boundaries of the homologous neighborhoods after the first, second, and third iterations, respectively. Protein sequences found after these iterations are colored correspondingly. Red, green, and blue numerals indicate the PROFILEs – the “centers of mass” of the sequences found in the above neighborhoods. This visualization does not define whether the search attains convergence after the third iteration or not. (A) The scenario in which the query sequence remains in the found list of homologs after all three iterations. This scenario mirrors the instance of the “DNA-directed RNA polymerase subunit epsilon” showcased in Fig. 2, where convergence was achieved after the third iteration. (B) The scenario where the query sequence is in the found list of homologs after the first and second iterations, but is excluded from it after the third iteration. This scenario aligns with the example of the “Ribonuclease HII” in Fig. 2, where the search loses the “Ribonuclease HII” after the third iteration but persists (*without* the “Ribonuclease HII”) for the next five iterations.

Figure 4A shows that losing the top position in the list of found homologs by the query sequence is more common than retaining it, even after the second iteration. Within the nr database-2011, merely 36%, 14%, 9%, and 8% of the sequences managed to maintain their first-place status in the list after the second, third, fourth, and fifth iterations. This phenomenon amplifies notably over time: for the nr 2021 database, these are as low as 18%, 5%, 3%, and 2%. The trend is clear: this pattern will only exacerbate as the database size continues to expand in the future.

**Figure 4.**
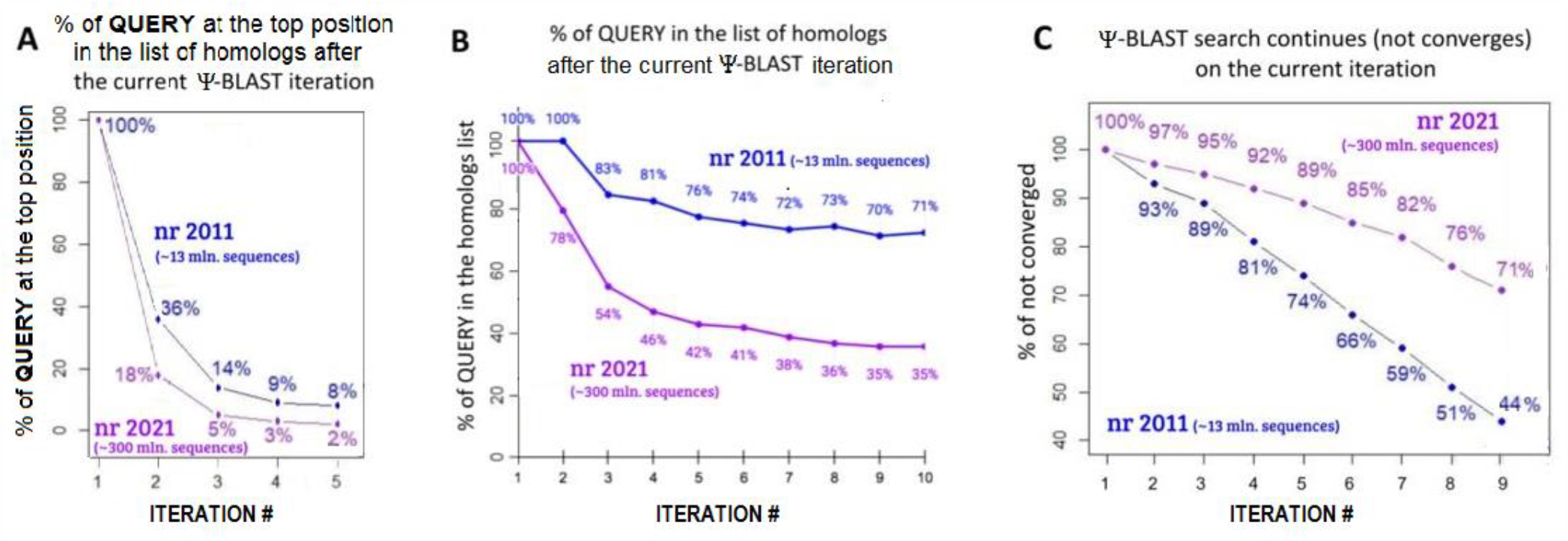
(A) The percentage of instances wherein the query sequence still occupies the top position in the list of homologs after the given iteration for the nr database of 2011 (blue) and 2021 (violet). For every percentage, the normalization was done by all 1000 Ψ-BLAST experiments. (B) The percentage of instances wherein the query sequence is present in the list of homologs after the current iteration for the nr database of 2011 (blue) and 2021 (violet). For each iteration, the % normalization was done by the number of the still ongoing - see panel C - Ψ-BLAST runs. (C) The percentage of the continued Ψ-BLAST searches vs. the iteration number for the nr database of 2011 (blue) and 2021 (violet). The % normalization was done by all 1000 Ψ-BLAST experiments.

However, even when the query sequence loses the top position, it can still influence the following Ψ-BLAST run as long as it is retained in the list of homologs. Figure 4B illustrates a gradual disappearance of query sequences from the lists of homologs yielded after subsequent iterations.

The worst occurs when the sequence completely disappears from the list of homologs defining the PROFILE (characterized by the symbol “>250” in Fig. 2), thereby no longer guiding the search. At the last completed Ψ-BLAST iteration (halted either by convergence or by the default limit of 10 iterations), the query sequence was present in the final list of homologs for the nr database-2021 in only 42% of cases, and absent in the rest 58%. Again, this negative effect becomes more pronounced over time: for the nr-database-2011, the values were “better”: 74% and 26% of cases, respectively.

Figure 4C shows that the percentage of still continued searches follows an almost linear decline with the iteration number. When considering the nr database-2011, half of the Ψ-BLAST searches attain convergence by the eighth iteration. In contrast, within the nr database-2021, only a quarter of the searches achieve convergence by the eighth iteration. It is evident that, with the database size constantly increasing over time, the challenge of achieving convergence intensifies. Notably, the dependence between the cases of unattained convergence and the iteration count (Fig.4C) provides a basis for estimating the number of iterations required for complete convergence of the Ψ-BLAST search across all sequences. According to this estimate, the convergence needs about 15 iterations for the nr database-2011 and about 30 iterations for the nr database-2021.

Thus, our results show the following: as the database size expands over time, Ψ-BLAST expedites the exclusion of the query sequence from the obtained list of homologs to this query sequence. This detracts the search for homologs from the original query sequence, and the Ψ-BLAST-found sequences can fail to be homologs of the query protein. Also, with the database expansion, the Ψ-BLAST convergence takes a longer time.

## Conclusions

In this study, we systematically explored the extremely popular Ψ-BLAST program with respect to its ability to keep focused on the query sequence throughout the process of searching for remotely homologous sequences. We aimed to figure out when the Ψ-BLAST (i) loses the query sequence from the top position in the obtained list of homologous sequences, and when (ii) it completely excludes the query sequence from this list during the iterative search. We discovered that merely after the second iteration the query sequence retains its top position only in 36% of cases for nr database-2011, and only in 18% for that of the year 2021. This reduces the efficacy of query sequence-centered search for the remote homologs. This is a pitfall of iterative Ψ-BLAST – a pitfall that arises just now. The situation which was tolerable 26 years ago when the Ψ-BLAST has been invented [Altschul et al., 1997] is now not as good, and it will exacerbate over time due to the database expansion.

In light of these circumstances, our recommendation is to monitor the position of the query sequence in the list of found homologs at each Ψ-BLAST iteration and use the disappearance of the query sequence from the BLAST list of homologs as a criterion to conclude the Ψ-BLAST search when the found sequences still remain related to the query sequence. Otherwise, the Ψ-BLAST-found sequences can fail to be homologs of the query protein.

## Acknowledgements

We acknowledge support from the Russian Science Foundation (grant no. 21-14-00268).

## Notes

### Competing Interest Statement

The authors have declared no competing interest.

